# Patterns of physical activity in hunter-gatherer children compared with US and UK children

**DOI:** 10.1101/2023.11.29.569171

**Authors:** Luke Kretschmer, Mark Dyble, Nikhil Chaudhary, David Bann, Gul Deniz Salali

**Affiliations:** Centre for Longitudinal Studies, Social Research Institute, UCL, London, UK; Department of Anthropology, University College London, 14 Taviton Street, London, UK; Department of Archaeology and Anthropology, University of Cambridge

**Keywords:** Physical Activity, Childhood, Hunter-Gatherers, Cross-Cultural, Evolution, Energetics

## Abstract

Sedentary lifestyles, mismatched with our active foraging history, contribute to escalating rates of non-communicable diseases. Contemporary hunter-gatherers appear to be highly active, but little is known about physical activity levels in hunter-gatherer children. We analysed 150 days of accelerometer data from 51 BaYaka hunter-gatherer children (aged 3-18) in the Republic of Congo, comparing it with British and American children (MCS and NHANES). BaYaka children were highly active, engaging in over 3 hours of moderate-to-vigorous physical activity (MVPA) daily, surpassing British adolescents by over 70 minutes. In US children activity declined with age; while in BaYaka children activity increased with age, unaffected by gender. Reflecting their foraging lifestyle, activity patterns varied within and between days, yet all children consistently rose with the sun. These findings highlight the impact of a foraging upbringing on children’s activity levels, providing a benchmark for understanding childhood physical activity and wellbeing.

## Introduction

With physical activity associated with increased lifespan and health span^1–3^ there is significant interest in examining the cultural factors driving increased activity. However, research that has attempted to do this has largely done so using culturally similar, highly sedentary, high-income populations.^4^ These populations represent a narrow fraction of human cultures, for whom daily calorie acquisition is decreasingly dependent on physical activity, with labour increasingly sedentary and foraging replaced with market-bought goods.^5^ Cross-cultural research on subsistence populations, including agriculturalists, horticulturalists, pastoralists, and hunter-gatherers, offer a valuable perspective as they reflect the diversity and history of humankind. Of these, hunter-gatherers may provide the best approximation of the physical activity patterns of the foraging lifestyle that dominated most of human evolutionary history.^6^ Notably, hunter-gatherer populations appear to have exceptionally low rates of non-communicable ‘diseases of modern life’, including low rates of obesity, type II diabetes, hypertension, and auto-immune disorders.^4, 6–8^ As such, any transferable insights may help inform meaningful interventions in post-industrial populations.

Studies of adults in subsistence populations reveal volumes of activity in excess of those seen in high-income populations.^9–11^ For example, a study of the Hadza hunter-gatherers of Tanzania observed daily amounts of moderate to vigorous physical activity (MVPA) to be in excess of weekly observations for US adults (135mins/day vs. 64mins/week)^12^ with similarly high volumes seen in other subsistence populations.^11, 13, 14^ However, a detailed understanding of the activity patterns of hunter-gatherer children is lacking. According to embodied capital theory, human childhood has evolved to allow necessary time to develop complex skills needed for the hunting and gathering niche.^15^ As such, hunter-gatherer childhoods are marked by learning of skills such as gathering wild plants, hunting animals, collecting honey and caterpillars, fishing, childcare and domestic activities.^16–18^ These skills often involve physically demanding activities such as climbing trees, using knives and machetes for digging and cutting, walking long distances in difficult terrain and carrying heavy loads including water and firewood. Although there is gendered division of labour, both boys and girls engage in laborious activities.^16, 19^ Hence, it is possible that unlike high-income populations where boys often undertake greater volumes of activity,^20^ hunter-gatherer boys and girls may engage in similar levels of activity. Forager children start practicing these skills from infancy, first by imitating and observing others, then by play and practice in mixed-aged autonomous children’s groups, often in the absence of adults.^16, 21^ Moreover, children also regularly act as caregivers, with children as young as 4 years old helping to care for babies, regularly holding and carrying them.^22, 23^

Given the low rates of physical activity among children in high-income populations and the importance of movement for lifelong health,^24–26^ it is important to establish comparative models of childhood physical activity. From the limited research detailing hunter-gatherer children’s activity levels, a study of space use in Baka children of Cameroon using waist-worn accelerometers observed children to undertake over 20,000 steps a day, with daily distances increasing with age.^27^ Another, based on observational data estimated an increase in moderate and vigorous intensity activity with age in Hadza children.^28^ There are, however, gaps in the current literature on hunter-gatherer children’s activity levels: 1) we lack direct comparisons of activity across cultures, 2) it is unclear whether boys and girls differ in their levels of activity as often observed in high-income populations, 3) the daily and hourly variation in activity levels are not well documented. In the absence of a regular routine of sedentary and active behaviours provided by classrooms, hunter-gatherer children may have more opportunity to vary in their activity within and between days.

In contrast to studies of hunter-gatherer populations, childhood physical activity is relatively well studied in high-income populations, with results that are largely replicated across populations.^20, 29^ Despite the well documented benefits of physical activity for healthy childhood development,^3, 30^ up to 80% of children globally fail to meet recommended physical activity guidelines.^31^ Research frequently observes that children tend to engage in diminishing volumes of physical activity with age,^32^ with reductions in activity notable following the start of formal schooling^33–36^ and following puberty,^36, 37^ with boys commonly undertaking more physical activity than girls.^38, 39^ Adding further relevance to childhood behaviours, the volumes of activity an individual engages in as they exit adolescence often predicts their volume of activity throughout adulthood.^40^

To address the paucity of data on physical activity among hunter-gatherer children, we examined patterns of physical activity among BaYaka children from the Republic of Congo using wrist-worn triaxial accelerometers. These devices allowed for examination of sedentary, light and moderate to vigorous intensities of activity, as well as the mean acceleration (presented as milligravitational units Euclidean norm minus one – mg ENMO). We additionally assessed differences in the sleep window (time of onset and wake-up) to examine differences in length of waking days. We compared patterns of activity between BaYaka children, with two nationally representative studies from high income countries, the American National Health And Nutrition Examination Survey (NHANES) and the British Millennium Cohort Study (MCS). We predict higher levels of physical activity, reduced gender difference in activity levels, and greater variability in daily activity levels in hunter-gatherer children compared to the children in the UK and the US.

## Results

### BaYaka children undertake three times more activity than WHO recommendations

Compared to the sampled high-income populations, BaYaka children were exceptionally active. Across the sample of 51 children (29 boys), the average day included over three hours of MVPA (Boys: 206min/day, Girls: 192min/day), a volume triple the WHO recommendations for children (Table S 1).^3^ The average 24-hour period included a further three hours of light intensity activity, with 17 hours a day spent sedentary, over 9 hours of which was wakeful. Together, this resulted in a mean acceleration of 47 mg ENMO (Boys: 48mg; Girls: 45mg; Table S 1). Compared to the average physical activity level of the British 14-year-olds in the UK MCS dataset, amongst the five 14-year-old BaYaka the mean acceleration was 16mg higher, with volumes of MVPA 65% greater (Figure 1 & 2).

**Figure 1:**
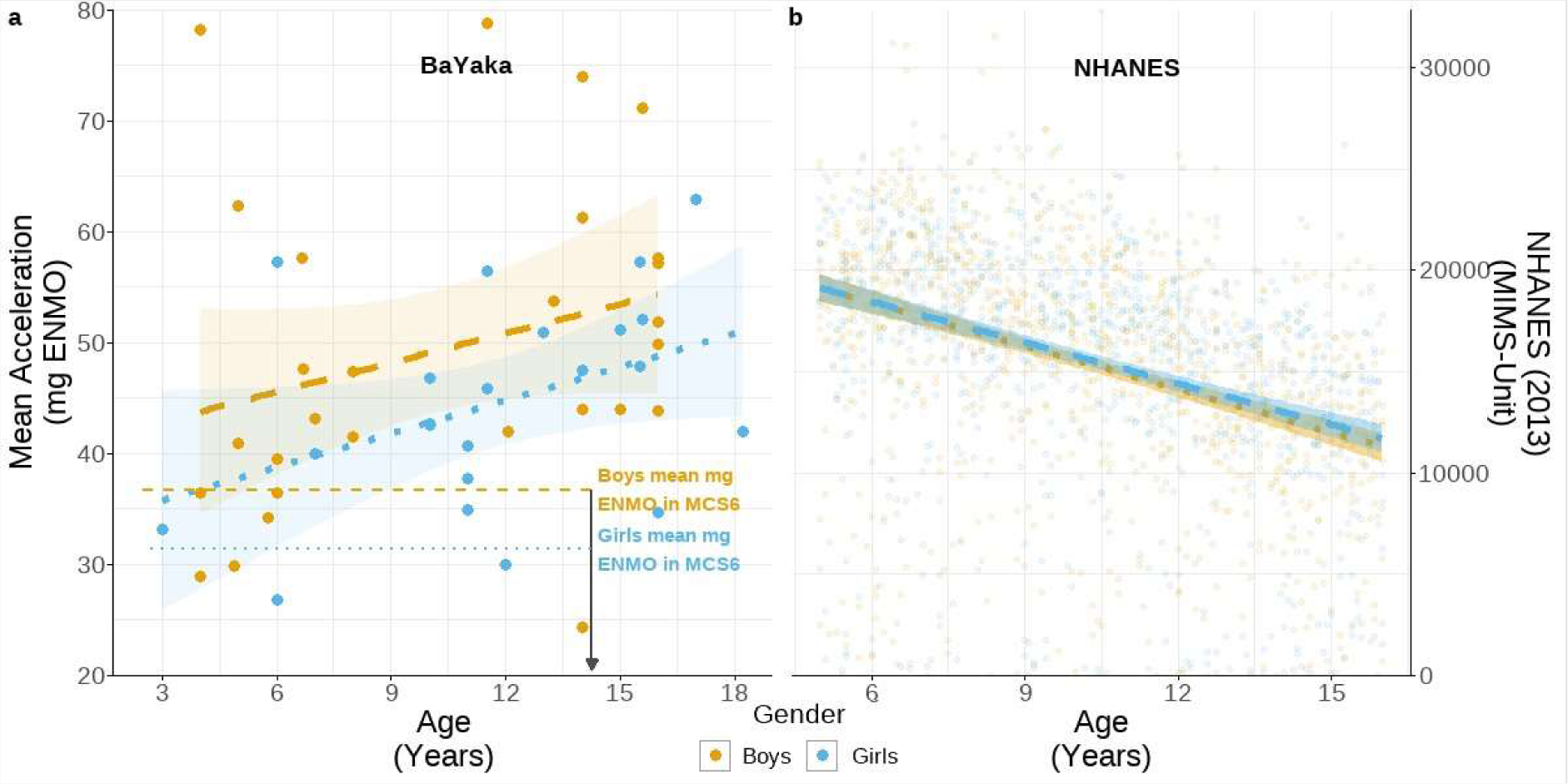
Mean acceleration amongst (a) BaYaka children as measued by mg ENMO and (b) US children in the 2013 NHANES study measured in MIMS-Unit. For both samples, all individuals aged between 3 and 18 years are included. Orange dots represent boys, blue dots represent girls. The fitted trend lines represent the linear model estimates and 95% CI as outlined in Table S 2. Horizontal dashed (boys) and dotted (girls) in panel a represent mean volumes observed in the Millennium Cohort Study (MCS6) where individuals were 14 years old. The Millennium Cohort Study processed physical activity in a manner consistent to that employed here, facilitating direct comparison among the 14-year-olds. Note that it is not possible to make direct comparisons of volumes of activity with the NHANES data due to the difference in the measurement unit.

### BaYaka children, regardless of gender, get more active with age unlike the children in the UK and the US

Overall activity increased with age in the BaYaka (+0.9 mg/year, CI: 0.1 – 1.6), with no meaningful difference between boys and girls (Girl: −4.5 mg, CI: −11.1 – 2.1; Table S 2). In both boys and girls activity was approximately 1mg higher for each additional year of age, maintaining a consistent trajectory between them (Figure 1, Table S 2). In contrast, the children in the US NHANES sample were more active in early childhood, and their activity declined with increasing age (Figure 1). Owing to differing means of reporting accelerations, internally calculated z-scored daily accelerations are used to assess differences between American and BaYaka children. Mean accelerations in the NHANES sample decreased with age by 0.51 standard deviations for each standard deviation of age (CI: −0.75 - −0.27), while BaYaka accelerations increased by 0.22 standard deviations for standard deviation of age (CI: −0.01 – 0.46). (Table S 3)

Separating overall activity in the BaYaka into the standard activity thresholds, the increase in mean acceleration was driven by greater volumes of moderate to vigorous intensity activities which in turn were offset by a decrease in sedentary activity (Figure 2). Volumes of MVPA was similar for both boys and girls (Girls: −24.1 min/day, CI: - 56.1 – 8.0), increasing by 8 minutes per day for each additional year of age (CI: 4.2 – 11.7). Offsetting this, volumes of sedentary behaviour decreased by an estimated 10 minutes per day for each additional year of age (CI: −14.8 - −3.4) (Table S 2), with no meaningful difference between boys and girls (Girls: +7.6 min/day, CI: −34.8 – 49.9). Of this, volumes of wakeful sedentary behaviour appear to be more stable with age, (--5 min/day CI: −13.4 – 2.7) indicating that differences in sedentary activity with age were also driven by changes in the sleep phase; the time between sleep onset and wake-up. As with sedentary behaviours, there were minimal changes to the estimated volumes of light intensity activity with age (+2.5 min/day, CI: 0.2 – 4.8).

**Figure 2:**
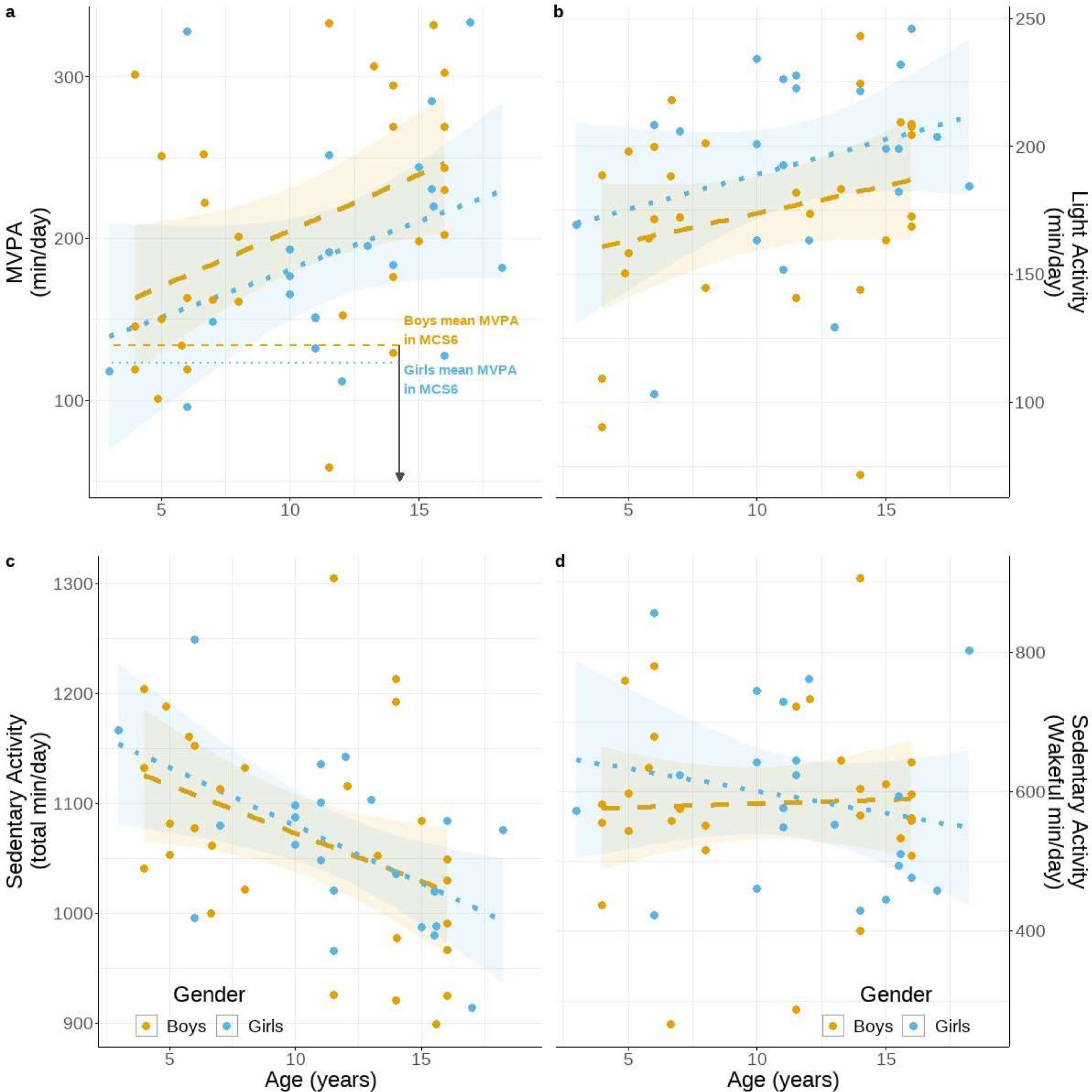
Mean volumes of activity per day by age. Boys are shown in orange and girls in blue (a) volume of moderate-to-vigorous intensity physical activity (mg ENMO > 100 mg) (b) volume of light intensity activity (mg ENMO: 50-99 mg), (c) volume of sedentary activity including sleep (mg ENMO < 50 mg), (d) volume of wakeful sedentary activity (mg ENMO < 50 mg). In panel a, the horizontal dashed (boys) and dotted (girls) represent mean volumes observed in the Millennium Cohort Study (MCS6) where individuals were 14 years old. The Millennium Cohort Study processed physical activity in a manner consistent to that employed here, facilitating direct comparison. Only values for MVPA and mg ENMO have been made available from MCS6. The volume of MVPA here includes all minutes above the threshold of 100 mg ENMO. The fitted trend lines represent the linear model estimates and 95% CI as outlined in Table S 2.

### Older children stay up later, but everyone wakes up together

Interestingly, BaYaka children had little variation in their wake-up time (+4min/year, CI: −5 – 13), but their time of sleep onset was later with increasing age (+10min/year, CI: 2 – 19) (Figure 3), with no difference between boys and girls (Table S 4). The reduced time in bed may partially underlie the reduction in sedentary behaviour with age, with sleep onset beginning 10 minutes later per year of age, which contributes to the decrease of 10 minutes a day of sedentary activity (Table S 2).

**Figure 3:**
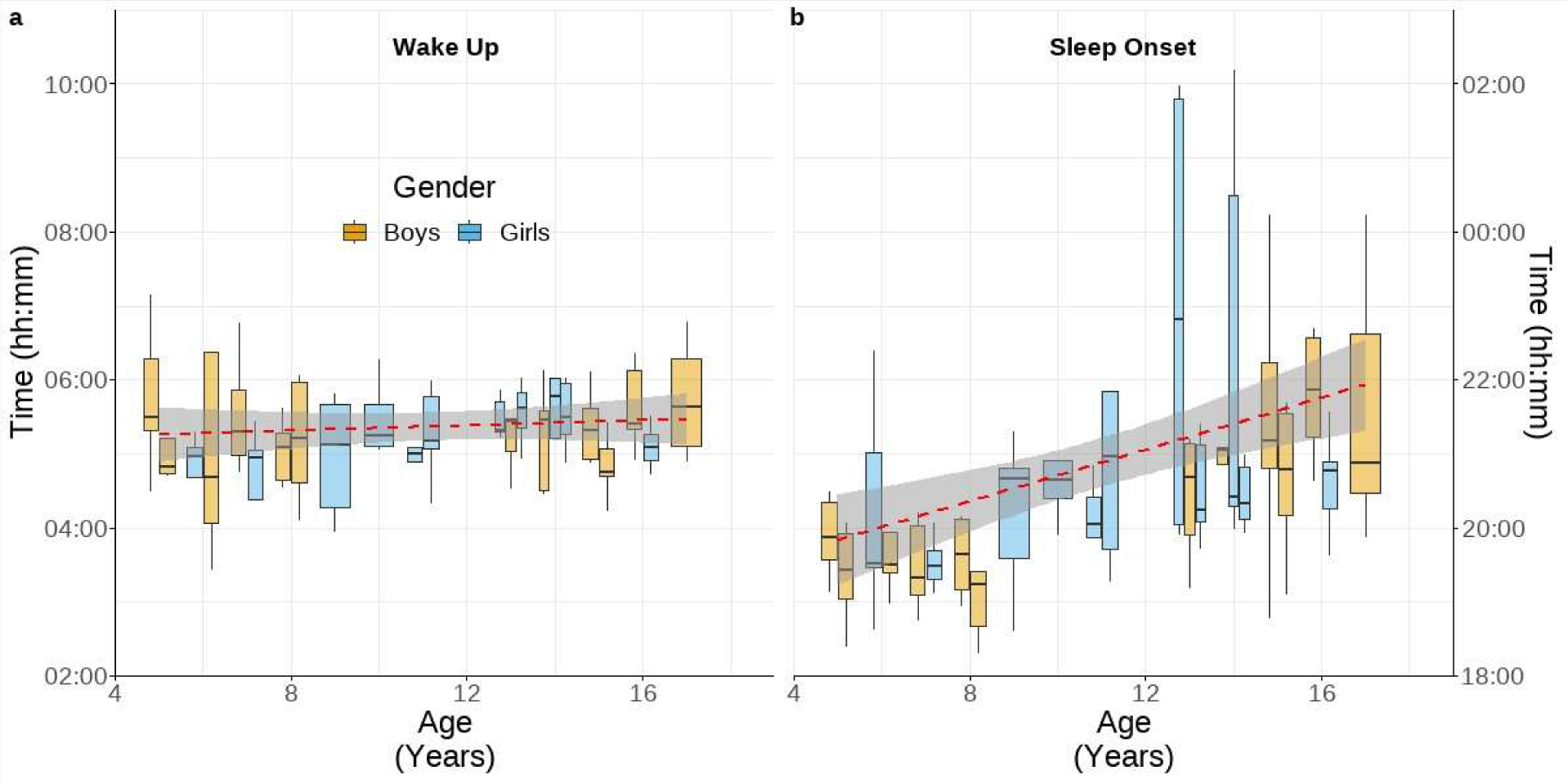
Device estimated (a) wake up and (b) sleep onset times across individuals. Dashed red line represents the estimate from linear model. Y-axes are matched in length to 9 hours each to aid interpretation. Sleep is classified by GGIR as any block of 10 minutes, in which the average movement of the device is less than 5° in any axis over each 5 second epoch. The algorithm defines the longest sustained inactive bout within each day as sleep, allowing for the sleep bout to be disrupted for up to one hour before. From this the estimated onset and end of sleep windows can be calculated as the final and first timepoint in each day in which an individual was not in a device defined bout of sleep

### Diurnal activity patterns reflect children’s foraging activities

BaYaka children’s activity patterns follow a clear diurnal pattern during the fishing season (Figure 4). Children were at their least active overnight, between 21:00 and 06:00, during which the average intensity of acceleration was below 10mg Transitioning from sleep, individuals rose to modest levels of activity in the early morning (06:00 to 09:00: 40mg) before being at their most active in the late morning (09:00 to 12:00: 80mg). During this most active period, children were twice as active as the previous time window. Throughout the afternoon (12:00 to 18:00) activity remained high and was stable at around 65mg. Following sunset, activity levels declined into the sleep window, with a mean acceleration of 42mg between 18:00 and 21:00. The two peaks in activity, one during early morning and the other late afternoon, likely reflect the walks children undertook when going into the forest for fishing and returning to the campsite after foraging (Figure 4). While children in the NHANES sample also undertook most of their activity during the day, compared to BaYaka, US children appeared to vary more in the times at which they woke-up, and remained active further into the night (Figure S 1 & Table S 6).

**Figure 4:**
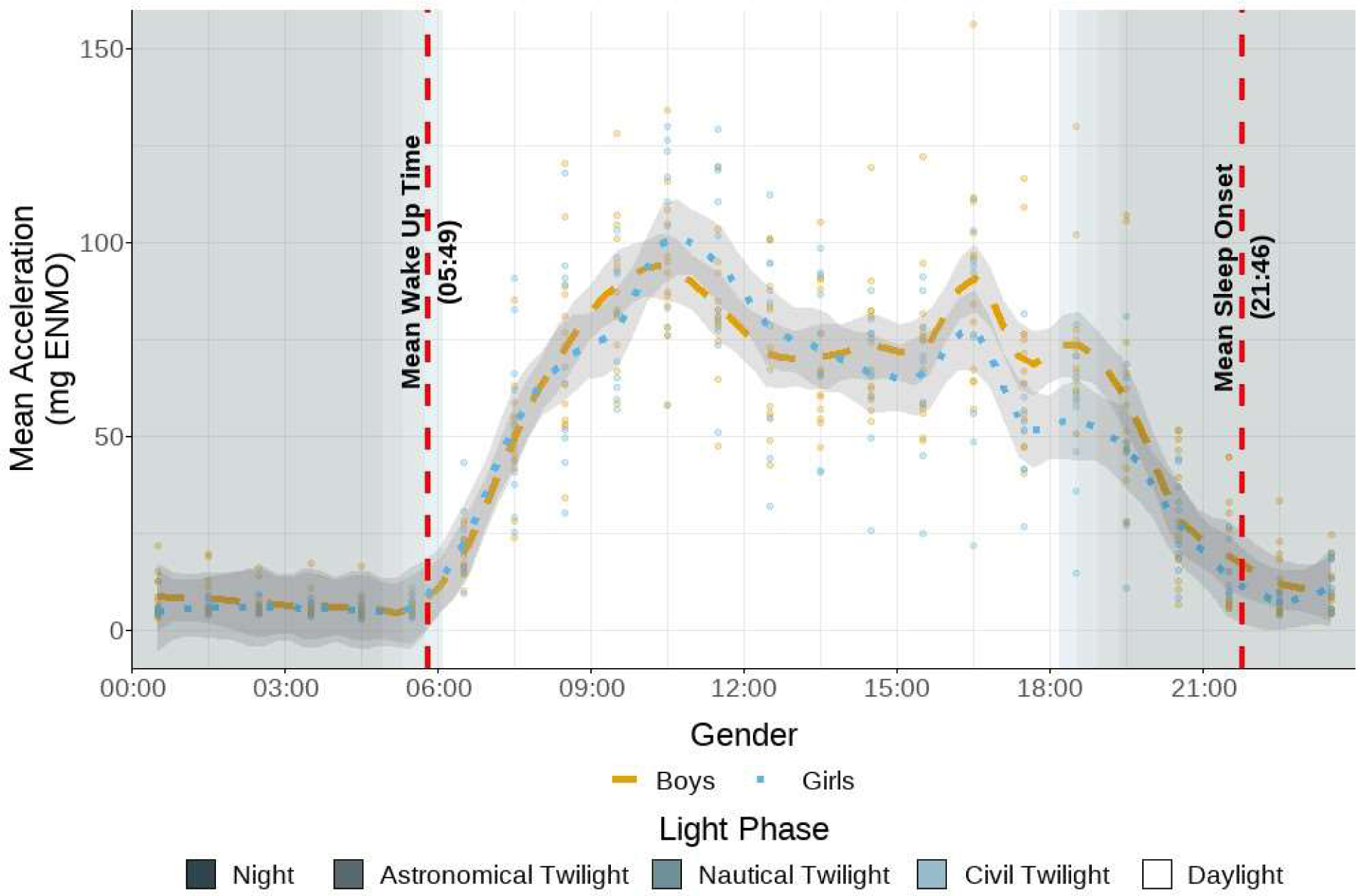
Distribution of physical activity throughout the day. For each individual in the 2022 data, their mean acceleration at each hour averaged across the 5 days of recording is presented as a single point, with colour corresponding to gender. The mean acceleration was calculated for each hour interval and presented at the half hour (i.e., the mean acceleration between 09:00 and 10:00 is presented at 09:30). Presented light phases were taken from the nearest town on the first day of recording. The dashed horizontal lines represent the mean time of wake (05:49) and sleep onset (21:46) across the sample presented in 24 hour hh:mm format.

### Activity levels in BaYaka children were more varied between days compared to the US children

Compared to US children in the NHANES dataset, BaYaka children exhibited more variation in how much activity they undertook each day. Using intercept only mixed effect models on standardised mean accelerations amongst BaYaka children with five days of recording (N=23) to match the length of recording in NHANES, a lower intra-class correlation (ICC) (0.50) was observed for BaYaka children than US children (0.66) (Table S 7). ICC measures variability within a group (here an individual’s ID) between 0 and 1 with scores closer to 0 indicating greater within-individual variation.

## Discussion

The BaYaka children included here were highly active, engaging in over three hours of MVPA per day on average, a figure three times higher than the WHO recommendations for children.^30^ Amongst the BaYaka, older children were more active than younger children, with greater mean accelerations driven by more time in MVPA (+8min/day for each year of age) exchanged with some sedentary time (−10min/day for each year of age). This is in stark contrast to children in the US sample presented here and other high-income populations where volumes of physical activity commonly peak in early childhood (ages 5 to 6) and decline from then, until reaching a low plateau in adulthood.^40^ Observational data amongst Hadza children also suggests an increase in MVPA with age,^28^ but using an objective measure here we observed BaYaka adolescents to engage in over 90 minutes more MVPA per day than British adolescents. Unlike those in the British cohort, no difference in activity was observed between BaYaka boys and girls, and this did not change with age. While adolescent BaYaka went to bed later, all children appeared to wake up together in time with sunrise. Moreover, BaYaka children showed more daily variation in their activity levels compared to the children in the US with their hourly activity patterns reflecting their foraging lifestyle. These results highlight the role of subsistence mode and potentially highlight the negative effect of formal schooling on children’s activity levels.

High levels of activity were observed from a young age, reinforcing observations that BaYaka children start practicing physically demanding foraging skills from early childhood.^16, 18^ Early childhood in hunter-gatherer and mixed foraging populations is marked by significant playtime in which children practice foraging, improve skills, and contribute to the family economy, either directly through food acquisition or indirectly through domestic work that frees other individuals to forage.^16, 23, 41^ Forager playgroups are critical for skill acquisition, often mimicking adult activities. For example, during 2022 fieldwork, we observed children aged 3-12 practicing dam fishing independently. This play and practice typically peak at age five for BaYaka children, then gradually shift towards work as they approach adolescence.^16, 41^ Activity increasing across childhood and adolescence mirrors the embodied capital theory in which a child’s skill, efficiency and the range of goods targeted increases as they grow.^15, 42^ Conversely, post-industrial populations see decreased activity over the same age span.^36^ A core difference between these children is in the behaviours they are increasingly modelling with age; for hunter-gatherer’s the archetypal adult is highly active, while parents in post-industrial populations are often less active than their children.^19, 43^ With the association between adolescent and adult activity, efforts to address adolescent activity may have lasting effects throughout adulthood.

BaYaka hunter-gatherers exhibit gendered division of labour with women targeting gatherable goods while men prioritise hunting and collecting honey. This division is reflected in children’s behaviour through their activities and play.^41^ Despite the gendered division of labour and differences in timing of play to work transition, activity was similar at all ages between BaYaka boys and girls. This contrast with the MCS study included here and other studies in industrialised populations, in which boys often engage in more physical activity.^33, 35, 38, 39, 44–49^ This highlights that the division in activity between boys and girls is likely of cultural rather than biological origin.

The 2022 fieldwork was done in dry season when BaYaka move to deeper parts of the forest and establish camps close to river streams to engage in fishing.^50^ During our fieldwork, BaYaka children frequently practiced dam fishing which involves making dams in the streams, bailing out the water, before killing the fish with machetes. The 2018 fieldwork occurred in July-August, a period when children engaged in treks into the forest to forage for caterpillars. Despite the seasonal differences in foraging activities, our results showed that the required activity levels for engaging in those are similar (Table S 2). Our hourly analysis also highlighted the diurnal trends in BaYaka children’s foraging which mimics adult foraging. The day was active with peaks of activity in late morning and early evening, coinciding with extended periods of walking to and from the forest.

The observations in the BaYaka and the sampled high-income populations suggest that formal schooling may promote inactivity in children, limiting their autonomy by mandating long periods of sedentary activity.^36, 51–54^ This schooling structure might contribute to increasing mental health problems and decreased child happiness observed in high-income populations.^54^ Without formal classrooms, BaYaka children choose their activities, with no imposed sedentary behaviour. Their daily activities, unlike American children’s regimented school schedules, are more variable and reflect an ecology of autonomous play, foraging, and rest. The structured, sedentary periods may impact children’s health and wellbeing, including adiposity increases.^55^ Implementing BaYaka perspectives, like breaking up prolonged bouts of sedentary behaviour as employed in forest-school and Udeskole (outdoor-school) programs has been observed to increase overall activity.^56, 57^

A teenage chronotype has been proposed to underlie later sleep onsets and wake up times amongst adolescents.^58^ Similar to studies in post-industrial populations, adolescent BaYaka did enter the sleep phase at a later time, possibly reflecting a reduction in sleep requirements with age.^59, 60^ However, there was little variation in wake-up times. For both, light is likely important; in the evening campfires allow the extension of wakeful activity safely into the night^61^ but does so without the blue wavelength light associated with disrupted sleep in industrialised populations.^62^ Then in the morning, with little ability to completely exclude light, daylight appears to be the entraining factor for waking up, as has previously been observed in Hadza adults.^63^

Highlighting modern mismatches in childhood physical activity, BaYaka children were highly active, engaging in over three times the WHO recommendations for physical activity. In further contrast to high-income populations, in a population that better embodies the ecology that humans evolved in, physical activity was more variable, but increased with age for boys and girls. In using cross-cultural studies to better understand the implications of low childhood activity on health, wellbeing and development, the use of highly active hunter-gatherer populations may provide a better reference point than another broadly inactive population.

## Methods

### Study population

The BaYaka are a group of hunter-gatherers residing in the rainforests of the Congo Basin. The Mbendjele are a subgroup of this population who speak Mbendjele.^64^ The BaYaka live in multifamily camps, consisting of 10-60 individuals,^65^ and move campsites based on the availability of forest products and trading opportunities throughout the year. The mode of subsistence is largely dependent on forest products: with food obtained by hunting, fishing and gathering wild products such as yams, caterpillars and honey depending on the season. This can be supplemented by trade with farmers and passing traders, with whom forest products are exchanged for market goods and cultivated foods.^66^

### Fieldwork and data collection

The BaYaka’s physical activity data was collected during fieldworks in 2018 and 2022 in the Likouala region of Congo’s Ndoki swamp forest. The 2018 fieldwork took place in July and August, when the main subsistence activities involved caterpillar and honey collecting, hunting, collecting wild yams and plants and occasional fishing. The 2022 fieldwork took place during fishing season in February when BaYaka move to deeper parts of the forest (leaving their camps by mud roads) and establish camps close to river streams to engage in fishing.

Between the two fieldtrips, 71 children aged up to 19 years wore a GENEActiv accelerometer device on their non-dominant arm. In 2018 46 children (24 boys) wore the device for up to two consecutive days. In 2022, 24 children (12 Boys) wore the device for 5 consecutive days. Children were excluded from the present analysis if they were aged under 3 years (n=6), if their accelerometery data was corrupted during recording (n=1) or if they did not have at least one complete day (midnight-midnight) of recording (n = 13). Together this left a sample of 51 children (2018 = 28, Total Boys = 29) totalling 150 days of activity recording (Table S 1).

Prior to distribution devices were set to run at 100hz, with participants instructed on how to wear the device, and that they should wear the device continuously throughout the recording. Age estimates for participants were established using a Bayesian method using relative age ranks based on those with known ages.^67^

The research and fieldworks were approved by the Ethics Committee of University College London (UCL Ethics codes 3086/003), and the methods were carried out in accordance with the approved guidelines. Informed consent was obtained from all participants and research permission granted by the Republic of Congo’s Ministry of Scientific Research.

### Physical activity metrics

Accelerometer recordings were processed using the R package GGIR.^68, 69^ The maximum recording length was set to 7 days from the start of recording. While no restriction on excluding the first or last day of study was stipulated, this was included de facto by excluding all incomplete days. Data was calculated in 24-hour windows from midnight to midnight, based on West Africa time (UTC +1 hour).

Acceleration is calculated as milligravitational units, minus one g to account for gravity (mg ENMO). Accelerations are presented as a continuous measure across the recording (mean mg ENMO) and subset into standard thresholds (Sedentary ≤ 50mg < Light ≤ 100mg < Moderate ≤ 400mg < Vigorous) consistent with previous research.^70, 71^ MVPA is a sum of all the minutes in moderate or vigorous intensities. Thresholds are presented as an unbouted measure, summing all time per day within an intensity threshold. Physical activity data for all individuals was collated for each individual at a daily level (1 entry per person per day of recording) and an hourly level (1 entry per person per hour of the day).

Sleep was detected by GGIR as periods of time in which the device was in a sustained, inactive state.^72^ This is classified as any block of 10 minutes, in which the average movement of the device is less than 5° in any axis over each 5 second epoch. The algorithm defines the longest sustained inactive bout within each day as sleep, allowing for the sleep bout to be broken for up to one hour before separating it into two bouts of sustained inactivity. From this the estimated onset and end of sleep windows can be calculated from the final and first timepoint in each day in which an individual was not in a device defined bout of sleep.

### Comparison populations

We used two publicly available datasets from high-income populations that had also employed wrist-worn accelerometers. One was the National Health and Nutrition Examination Survey (NHANES) and the other was the 6^th^ sweep of the UK Millennium Cohort Study (MCS). NHANES is a repeated cross-sectional study conducted across the United States and designed to be broadly representative of the US population, with some oversampling of minority groups to ensure statistical power when subset.^73^ The study was conducted in 2013 to 2014 with individuals aged 3 years or older wearing a wrist-worn triaxial accelerometer (Actigraph GT3X+) for 7 continuous days. We used a subset of this data to match the BaYaka sample. The included NHANES sample contained 2391 children (1200 boys) aged between 5 and 18 years (Table S 9). The mean age was 11 years 8 months and was similar between boys and girls (boys: 10.60 years; girls: 10.76 years). Of the individuals included, 25% identified as non-Hispanic White, 25% and non-Hispanic Black, 25% as Mexican American, 10% as other Hispanic and 15% as ‘other’ ethnicity. The mean MIMS-Unit (Monitor Independent Motion Sensing) were 14,673 and were similar between boys and girls (boys: 14,534 MIMS; girls: 14,813 MIMS). Due to differences in output metrics, we did not compare the volumes of activity directly between NHANES and the BaYaka (See Table S 9). Instead, we used internally estimated z-scored average activity measures to examine variation across ages.

The MCS accelerometery data included 4,533 individuals (2,182 boys) aged 14 years (Table S 8).^74, 75^ Because the data was collected using the same device as the BaYaka data, we were able to make direct comparisons among the 14 year-olds. Data was processed using GGIR in the same manner as done with the BaYaka with average accelerations and time in MVPA calculated in the same manner and presented in the same units.

## Statistical analyses

### Modelling BaYaka activity

To model difference between ages and genders, linear mixed effect models were employed, using the lme4 package in R.^69, 76^ Outcome variables were either the mean acceleration across the day or the volume of time in a given threshold of activity. Models were adjusted for age and gender, both as additive effects, and with an interaction term to allow gender specific slopes for age. Additionally, year of recording was included in the models to account for potential seasonal differences between the two visits to the field. Individual ID was included as a random effect to adjust for clustering of values within an individual across the days of recording. The sample was restricted to include only days with a complete 24hrs of recording, and to individuals who had complete data for the included covariates. This left a sample of 150 days of activity shared between 51 individuals.

Mixed effect models were also used to analyse differences in activity across the day. Mean accelerations were averaged over 3-hour windows, for each day of recording for each child. The effect of time of day (in 3-hour windows) was modelled against the mean acceleration in that window, with the fixed covariates of age and gender included. To account for the nested clustering of results within days and within individuals, random effects of day and individual ID were included to reflect this nesting of groups. Because 2022 data included a longer period, we used this data for analysis of within day variation in activity. The reference in these models mirrors previous models for overall activity, with the reference time window being in the middle of the sleep window (midnight – 3am). Three-hour windows were used to improve the contrast between windows of time and reduce the number of pairwise comparisons.

To model differences in wake-up times and sleep-onset, data from the 2022 data collection was used. With recording conducted over the same week for all individuals, light conditions were the same for all individuals. For each of wake-up and sleep onset, the outcome of time of day is modelled against the effect of age and gender as fixed effects with the day of recording nested within each individual as a random effect.

### Modelling patterns of activity in the BaYaka and the US sample

To compare the BaYaka and NHANES study, the mean acceleration on each day of recording (measured as mg ENMO & MIMS Unit) was standardised using internally calculated z-scores. Z-scored mean accelerations were then employed in mixed effect linear models adjusted for age, gender and population as fixed effects, with the individual ID included as a random effect. To examine differences between populations in the slope with age, an interaction term was included between age and population.

## Supplemental Information

### Data Availability

Data is available on request. Details and access to the NHANES dataset can be found here (https://www.cdc.gov/nchs/nhanes/index.htm). Detail on the Millennium Cohort Study can be found here (https://cls.ucl.ac.uk/cls-studies/millennium-cohort-study/) and can be accessed via the UK data service.

### Code Availability

A markdown script detailing all included analyses is available at http://rpubs.com/LukeKretschmer/1116124

## Acknowledgements

Many thanks to all the Mbendjele BaYaka participants for their hospitality and participation; our BaYaka field assistants Nicolas Yuppe, and Dambo Gaston; and Selcen Kucukustel, Sarai Keestra, Inez Derkx and Gaurav Sikka for their help in data collection; and Dr Edmond Sylvestre MIABANGANA, Dr Laure Stella Ghoma Linguissi and Dr Guy Moussavou for their help with research permits. Thanks are also extended to Dr Megan Arnot for her help developing the included figures.

## Author Contributions

GDS conceived the project. GDS collected the 2022 data, GDS and NC collected the 2018 data. LK analysed the data and wrote the first draft under the supervision of GDS, DB and MD. All authors contributed to reviewing and editing of the manuscript.

## References

1. Warburton, D. E. R., & Bredin, S. S. D. (2017). Health benefits of physical activity: A systematic review of current systematic reviews. Current Opinion in Cardiology, 32(5), 541–556.

2. Ekelund, U., Tarp, J., Steene-Johannessen, J., Hansen, B. H., Jefferis, B., Fagerland, M. W., Whincup, P., Diaz, K. M., Hooker, S. P., Chernofsky, A., Larson, M. G., Spartano, N., Vasan, R. S., Dohrn, I. M., Hagströmer, M., Edwardson, C., Yates, T., Shiroma, E., Anderssen, S. A., & Lee, I. M. (2019). Dose-response associations between accelerometry measured physical activity and sedentary time and all cause mortality: Systematic review and harmonised meta-analysis. The BMJ, 366.

3. World Health Organization. (2020). WHO Guidelines On Physical Activity And Sedentary Behaviour.

4. Gurven, M. D., & Lieberman, D. E. (2020). WEIRD bodies: mismatch, medicine and missing diversity.

5. Brownson, R. C., Boehmer, T. K., & Luke, D. A. (2005). Declining rates of physical activity in the United States: what are the contributors? Annual Review of Public Health, 26, 421–443.

6. Pontzer, H., Wood, B. M., & Raichlen, D. A. (2018). Hunter-gatherers as models in public health. Obesity Reviews, 19, 24–35.

7. Madimenos, F. C., Snodgrass, J. J., Blackwell, A. D., Liebert, M. A., & Sugiyama, L. S. (2011). Physical activity in an indigenous Ecuadorian forager-horticulturalist population as measured using accelerometry. American Journal of Human Biology, 23(4), 488–497.

8. Gurven, M., Stieglitz, J., Trumble, B., Blackwell, A. D., Beheim, B., Davis, H., Hooper, P., & Kaplan, H. (2017). The Tsimane Health and Life History Project: Integrating anthropology and biomedicine. Evolutionary Anthropology: Issues, News, and Reviews, 26(2), 54–73.

9. Yamauchi, T., Sato, H., & Kawamura, K. (2000). Nutritional Status, Activity Pattern, And Dietary Intake Among The Baka Hunter-Gatherers In The Village Camps In Cameroon. African Study Monographs, 21(2), 67–82.

10. Ojiambo, R., Gibson, A. R., Konstabel, K., Lieberman, D. E., Speakman, J. R., Reilly, J. J., & Pitsiladis, Y. P. (2013). Free-living physical activity and energy expenditure of rural children and adolescents in the Nandi region of Kenya. Annals of Human Biology, 40(4), 318–323.

11. Sarma, M. S., Boyette, A. H., Lew-Levy, S., Miegakanda, V., Kilius, E., Samson, D. R., & Gettler, L. T. (2020). Sex differences in daily activity intensity and energy expenditure and their relationship to cortisol among <scp>BaYaka</scp> foragers from the Congo Basin. American Journal of Physical Anthropology, ajpa.24075.

12. Raichlen, D. A., Pontzer, H., Harris, J. A., Mabulla, A. Z. P., Marlowe, F. W., Josh Snodgrass, J., Eick, G., Colette Berbesque, J., Sancilio, A., & Wood, B. M. (2017). Physical activity patterns and biomarkers of cardiovascular disease risk in hunter-gatherers. American Journal of Human Biology, 29(2), e22919.

13. Gurven, M., Jaeggi, A. V., Kaplan, H., & Cummings, D. (2013). Physical Activity and Modernization among Bolivian Amerindians. PLoS ONE, 8(1), e55679.

14. Sayre, M. K., Pike, I. L., & Raichlen, D. A. (2019). High levels of objectively measured physical activity across adolescence and adulthood among the Pokot pastoralists of Kenya. American Journal of Human Biology, 31(1).

15. Kaplan, H., Lancaster, J., & Robson, A. (2003). Embodied Capital and the Evolutionary Economics of the Human Life Span. Population and Development Review, 29, 152–182.

16. Salali, G. D., Chaudhary, N., Bouer, J., Thompson, J., Vinicius, L., & Migliano, A. B. (2019). Development of social learning and play in BaYaka hunter-gatherers of Congo. Scientific Reports, 9(1), 11080.

17. Salali, G. D., Chaudhary, N., Thompson, J., Grace, O. M., van der Burgt, X. M., Dyble, M., Page, A. E., Smith, D., Lewis, J., Mace, R., Vinicius, L., & Migliano, A. B. (2016). Knowledge-Sharing Networks in Hunter-Gatherers and the Evolution of Cumulative Culture. Current Biology: CB, 26(18), 2516–2521.

18. Lew-Levy, S., Reckin, R., Lavi, N., Cristóbal-Azkarate, J., & Ellis-Davies, K. (2017). How Do Hunter-Gatherer Children Learn Subsistence Skills? Human Nature, 28(4), 367–394.

19. Pontzer, H., Raichlen, D. A., Wood, B. M., Emery Thompson, M., Racette, S. B., Mabulla, A. Z. P., & Marlowe, F. W. (2015). Energy expenditure and activity among Hadza hunter-gatherers. American Journal of Human Biology, 27(5), 628–637.

20. Kretschmer, L., Salali, G. D., Andersen, L. B., Hallal, P. C., Northstone, K., Sardinha, L. B., Dyble, M., Bann, D., Andersen, L. B., Anderssen, S., Cardon, G., Davey, R., Jago, R., Janz, K. F., Kriemler, S., Møller, N., Northstone, K., Pate, R., Puder, J. J., … International Children’s Accelerometry Database (ICAD) Collaborators. (2023). Gender differences in the distribution of children’s physical activity: evidence from nine countries. International Journal of Behavioral Nutrition and Physical Activity, 20(1), 103.

21. Boyette, A. H., & Hewlett, B. S. (2017). Autonomy, Equality, and Teaching among Aka Foragers and Ngandu Farmers of the Congo Basin. Human Nature, 28(3), 289–322.

22. Mesman, J., Minter, T., Angnged, A., Cissé, I. A. H., Salali, G. D., & Migliano, A. B. (2018). Universality Without Uniformity: A Culturally Inclusive Approach to Sensitive Responsiveness in Infant Caregiving. Child Development, 89(3), 837–850.

23. Page, A. E., Emmott, E. H., Dyble, M., Smith, D., Chaudhary, N., Viguier, S., & Migliano, A. B. (2021). Children are important too: juvenile playgroups and maternal childcare in a foraging population, the Agta. Philosophical Transactions of the Royal Society B: Biological Sciences, 376(1827), rstb.2020.0026.

24. Veldman, S. L. C., Chin A Paw, M. J. M., & Altenburg, T. M. (2021). Physical activity and prospective associations with indicators of health and development in children aged <5 years: a systematic review. International Journal of Behavioral Nutrition and Physical Activity, 18(1), 1–11.

25. Boreham, C., & Riddoch, C. (2001). The physical activity, fitness and health of children. Journal of Sports Sciences, 19(12), 915–929.

26. García-Hermoso, A., Ramírez-Vélez, R., García-Alonso, Y., Alonso-Martínez, A. M., & Izquierdo, M. (2020). Association of Cardiorespiratory Fitness Levels During Youth With Health Risk Later in Life: A Systematic Review and Meta-analysis. JAMA Pediatrics, 174(10), 952–960.

27. Hagino, I., & Yamauchi, T. (2014). Daily Physical Activity and Time-Space Using of Pygmy Hunter-Gatherers’ Children in Southeast Cameroon. In Dynamics of Learning in Neanderthals and Modern Humans Volume 2 (pp. 91–97). Springer Japan.

28. Froehle, A. W., Wells, G. K., Pollom, T. R., Mabulla, A. Z. P., Lew-Levy, S., & Crittenden, A. N. (2019). Physical activity and time budgets of Hadza forager children: Implications for self-provisioning and the ontogeny of the sexual division of labor. American Journal of Human Biology, 31(1), e23209.

29. Guthold, R., Stevens, G. A., Riley, L. M., & Bull, F. C. (2020). Global trends in insufficient physical activity among adolescents: a pooled analysis of 298 population-based surveys with 1·6 million participants. The Lancet Child and Adolescent Health, 4(1), 23–35. Scopus.

30. World Health Organisation. (2019). WHO guidelines on physical activity, sedentary behaviour and sleep for children under 5 years of age. WHO.

31. World Health Organisation. (2022). Global status report on physical activity 2022. https://www.who.int/publications-detail-redirect/9789240059153

32. Farooq, A., Martin, A., Janssen, X., Wilson, M. G., Gibson, A. M., Hughes, A., & Reilly, J. J. (2020). Longitudinal changes in moderate-to-vigorous-intensity physical activity in children and adolescents: A systematic review and meta-analysis. Obesity Reviews, 21(1), e12953.

33. Cooper, A. R., Goodman, A., Page, A. S., Sherar, L. B., Esliger, D. W., van Sluijs, E. M. F., Andersen, L. B., Anderssen, S., Cardon, G., Davey, R., Froberg, K., Hallal, P., Janz, K. F., Kordas, K., Kreimler, S., Pate, R. R., Puder, J. J., Reilly, J. J., Salmon, J., … Ekelund, U. (2015). Objectively measured physical activity and sedentary time in youth: The International children’s accelerometry database (ICAD). International Journal of Behavioral Nutrition and Physical Activity, 12(1), 1–10.

34. Aadland, E., Okely, A. D., Kristine, A., & Nilsen, O. (2022). Trajectories of physical activity and sedentary time in Norwegian children aged 3–9 years: a 5-year longitudinal study. International Journal of Behavioral Nutrition and Physical Activity 2022 19:1, 19(1), 1–13.

35. Steene-Johannessen, J., Hansen, B. H., Dalene, K. E., Kolle, E., Northstone, K., Møller, N. C., Grøntved, A., Wedderkopp, N., Kriemler, S., Page, A. S., Puder, J. J., Reilly, J. J., Sardinha, L. B., Van Sluijs, E. M. F., Andersen, L. B., Van Der Ploeg, H., Ahrens, W., Flexeder, C., Standl, M., … Van Sluijs, E. M. F. (2020). Variations in accelerometry measured physical activity and sedentary time across Europe-harmonized analyses of 47,497 children and adolescents. International Journal of Behavioral Nutrition and Physical Activity, 17(1), 1–14.

36. Reilly, J. J. (2016). When does it all go wrong? Longitudinal studies of changes in moderate-to-vigorous-intensity physical activity across childhood and adolescence. Journal of Exercise Science & Fitness, 14(1), 1–6.

37. Corder, K., Sharp, S. J., Atkin, A. J., Griffin, S. J., Jones, A. P., Ekelund, U., & Van Sluijs, E. M. F. (2015). Change in objectively measured physical activity during the transition to adolescence. British Journal of Sports Medicine, 49(11), 730– 736.

38. Bauman, A. E., Reis, R. S., Sallis, J. F., Wells, J. C., Loos, R. J. F., Martin, B. W., Alkandari, J. R., Andersen, L. B., Blair, S. N., Brownson, R. C., Bull, F. C., Craig, C. L., Ekelund, U., Goenka, S., Guthold, R., Hallal, P. C., Haskell, W. L., Heath, G. W., Inoue, S., … Sarmiento, O. L. (2012). Correlates of physical activity: Why are some people physically active and others not? The Lancet, 380(9838), 258–271.

39. Trost, S. G., Pate, R. R., Sallis, J. F., Freedson, P. S., Taylor, W. C., Dowda, M., & Sirard, J. (2002). Age and gender differences in objectively measured physical activity in youth. In Med. Sci. Sports Exerc (Vol. 34, Issue 2, pp. 350– 355). http://www.acsm-msse.org

40. Varma, V. R., Dey, D., Leroux, A., Di, J., Urbanek, J., Xiao, L., & Zipunnikov, V. (2017). Re-evaluating the effect of age on physical activity over the lifespan. Preventive Medicine, 101, 102–108.

41. Lew-Levy, S., Reckin, R., Kissler, S. M., Pretelli, I., Boyette, A. H., Crittenden, A. N., Hagen, R. V., Haas, R., Kramer, K. L., Koster, J., O’Brien, M. J., Sonoda, K., Surovell, T. A., Stieglitz, J., Tucker, B., Lavi, N., Ellis-Davies, K., & Davis, H. E. (2022). Socioecology shapes child and adolescent time allocation in twelve hunter-gatherer and mixed-subsistence forager societies. Scientific Reports, 12(1), 8054.

42. Koster, J., McElreath, R., Hill, K., Yu, D., Shepard, G., Van Vliet, N., Gurven, M., Trumble, B., Bird, R. B., Bird, D., Codding, B., Coad, L., Pacheco-Cobos, L., Winterhalder, B., Lupo, K., Schmitt, D., Sillitoe, P., Franzen, M., Alvard, M., … Ross, C. (2020). The life history of human foraging: Cross-cultural and individual variation. Science Advances, 6(26), 9070–9094.

43. Belcher, B. R., Wolff-Hughes, D. L., Dooley, E. E., Staudenmayer, J., Berrigan, D., Eberhardt, M. S., & Troiano, R. P. (2021). US Population-referenced Percentiles for Wrist-Worn Accelerometer-derived Activity. Medicine and Science in Sports and Exercise, 53(11), 2455–2464.

44. Corder, K., Sharp, S. J., Atkin, A. J., Andersen, L. B., Cardon, G., Page, A., Davey, R., Grøntved, A., Hallal, P. C., Janz, K. F., Kordas, K., Kriemler, S., Puder, J. J., Sardinha, L. B., Ekelund, U., van Sluijs, E. M. F., Cardon, G., Cooper, A., Puder, J. J., … Timperio, A. (2016). Age-related patterns of vigorous-intensity physical activity in youth: The International Children’s Accelerometry Database. Preventive Medicine Reports, 4, 17–22.

45. Dias, K. I., White, J., Jago, R., Cardon, G., Davey, R., Janz, K. F., Pate, R. R., Puder, J. J., Reilly, J. J., & Kipping, R. (2019). International Comparison of the Levels and Potential Correlates of Objectively Measured Sedentary Time and Physical Activity among Three-to-Four-Year-Old Children. International Journal of Environmental Research and Public Health 2019, Vol. 16, Page 1929, 16(11), 1929.

46. Van Ekris, E., Wijndaele, K., Altenburg, T. M., Atkin, A. J., Twisk, J., Andersen, L. B., Janz, K. F., Froberg, K., Northstone, K., Page, A. S., Sardinha, L. B., Van Sluijs, E. M. F., Chinapaw, M., Andersen, L. B., Anderssen, S., Atkin, A. J., Cardon, G., Davey, R., Ekelund, U., … Van Sluijs, E. M. F. (2020). Tracking of total sedentary time and sedentary patterns in youth: A pooled analysis using the International Children’s Accelerometry Database (ICAD). International Journal of Behavioral Nutrition and Physical Activity, 17(1), 1–10.

47. Kwon, S., Janz, K. F., Cooper, A., Ekelund, U., Esliger, D., Griew, P., Judge, K., Ness, A., Riddoch, C., Salmon, J., & Sherar, L. (2012). Tracking of accelerometry-measured physical activity during childhood: ICAD pooled analysis. International Journal of Behavioral Nutrition and Physical Activity, 9(1), 1–8.

48. Tarp, J., Child, A., White, T., Westgate, K., Bugge, A., Grøntved, A., Wedderkopp, N., Andersen, L. B., Cardon, G., Davey, R., Janz, K. F., Kriemler, S., Northstone, K., Page, A. S., Puder, J. J., Reilly, J. J., Sardinha, L. B., van Sluijs, E. M. F., Ekelund, U., … Brage, S. (2018). Physical activity intensity, bout-duration, and cardiometabolic risk markers in children and adolescents. International Journal of Obesity 2018 42:9, 42(9), 1639–1650.

49. Konstabel, K., Veidebaum, T., Verbestel, V., Moreno, L. A., Bammann, K., Tornaritis, M., Eiben, G., Molnár, D., Siani, A., Sprengeler, O., Wirsik, N., Ahrens, W., & Pitsiladis, Y. (2014). Objectively measured physical activity in European children: the IDEFICS study. International Journal of Obesity 2014 38:2, 38(2), S135–S143.

50. Jang, H., & Boyette, A. H. (2021). Observations of Cooperative Pond Fishing by the Bayaka and Bantu People in the Flooded Forest of the Northern Republic of Congo. African Study Monographs, 41(2), 1–16.

51. Chaudhary, N., & Swanepoel, A. (2023). Editorial Perspective: What can we learn from hunter-gatherers about children’s mental health? An evolutionary perspective. Journal of Child Psychology and Psychiatry, n/a(n/a).

52. Arundell, L., Hinkley, T., Veitch, J., & Salmon, J. (2015). Contribution of the After-School Period to Children’s Daily Participation in Physical Activity and Sedentary Behaviours. PLOS ONE, 10(10), e0140132.

53. McLellan, G., Arthur, R., Donnelly, S., & Buchan, D. S. (2020). Segmented sedentary time and physical activity patterns throughout the week from wrist-worn ActiGraph GT3X+ accelerometers among children 7–12 years old. Journal of Sport and Health Science, 9(2), 179–188.

54. Chaudhary, N., & Salali, G. D. (2022). Hunter-Gatherers, Mismatch and Mental Disorder. In P. St John-Smith & R. Abed (Eds.), Evolutionary Psychiatry: Current Perspectives on Evolution and Mental Health (pp. 64–83). Cambridge University Press.

55. Wijndaele, K., White, T., Andersen, L. B., Bugge, A., Kolle, E., Northstone, K., Wedderkopp, N., Ried-Larsen, M., Kriemler, S., Page, A. S., Puder, J. J., Reilly, J. J., Sardinha, L. B., Van Sluijs, E. M. F., Sharp, S. J., Brage, S., Ekelund, U., Andersen, L. B., Atkin, A. J., … Van Sluijs, E. M. F. (2019). Substituting prolonged sedentary time and cardiovascular risk in children and youth: A meta-analysis within the International Children’s Accelerometry database (ICAD). International Journal of Behavioral Nutrition and Physical Activity, 16(1), 1–10.

56. Mygind, E. (2016). Physical Activity during Learning Inside and Outside the Classroom. Health Behavior and Policy Review, 3(5), 455–467.

57. Dabaja, Z. F. (2022). The Forest School impact on children: reviewing two decades of research. Education 3-13, 50(5), 640–653.

58. Hagenauer, M. H., & Lee, T. M. (2013). Adolescent sleep patterns in humans and laboratory animals. Hormones and Behavior, 64(2), 270–279.

59. Williams, J. A., Zimmerman, F. J., & Bell, J. F. (2013). Norms and Trends of Sleep Time Among US Children and Adolescents. JAMA Pediatrics, 167(1), 55–60.

60. Paruthi, S., Brooks, L. J., D’Ambrosio, C., Hall, W. A., Kotagal, S., Lloyd, R. M., Malow, B. A., Maski, K., Nichols, C., Quan, S. F., Rosen, C. L., Troester, M. M., & Wise, M. S. (2016). Consensus Statement of the American Academy of Sleep Medicine on the Recommended Amount of Sleep for Healthy Children: Methodology and Discussion. Journal of Clinical Sleep Medicine: JCSM: Official Publication of the American Academy of Sleep Medicine, 12(11), 1549– 1561.

61. Dunbar, R. I. M., Gamble, C., & Gowlett, J. A. J. (2014). Fireside Chat: The Impact of Fire on Hominin Socioecology. In Lucy to Language: The Benchmark Papers. OUP Oxford.

62. Tähkämö, L., Partonen, T., & Pesonen, A.-K. (2019). Systematic review of light exposure impact on human circadian rhythm. Chronobiology International, 36(2), 151–170.

63. Samson, D. R., Crittenden, A. N., Mabulla, I. A., Mabulla, A. Z. P., & Nunn, C. L. (2017). Hadza sleep biology: Evidence for flexible sleep-wake patterns in hunter-gatherers. American Journal of Physical Anthropology, 162(3), 573– 582.

64. Lewis, J. (2015). Where goods are free but knowledge costs. Hunter Gatherer Research, 1(1).

65. Dyble, M., Salali, G. D., Chaudhary, N., Page, A., Smith, D., Thompson, J., Vinicius, L., Mace, R., & Migliano, A. B. (2015). Sex equality can explain the unique social structure of hunter-gatherer bands. Science, 348(6236), 796 LP – 798.

66. Knight, J. K., Salali, G. D., Sikka, G., Derkx, I., Keestra, S. M., & Chaudhary, N. (2021). Quantifying patterns of alcohol consumption and its effects on health and wellbeing among BaYaka hunter-gatherers: A mixed-methods cross-sectional study. PLOS ONE, 16(10), e0258384.

67. Diekmann, Y., Smith, D., Gerbault, P., Dyble, M., Page, A. E., Chaudhary, N., Migliano, A. B., & Thomas, M. G. (2017). Accurate age estimation in small-scale societies. Proceedings of the National Academy of Sciences of the United States of America, 114(31), 8205–8210.

68. Migueles, J. H., Rowlands, A. V., Huber, F., Sabia, S., & van Hees, V. T. (2019). GGIR: A Research Community–Driven Open Source R Package for Generating Physical Activity and Sleep Outcomes From Multi-Day Raw Accelerometer Data. Journal for the Measurement of Physical Behaviour, 2(3), 188–196.

69. R Core Team. (2020). R: A Language and Environment for Statistical Computing. R Foundation for Statistical Computing. http://www.r-project.org/

70. Kontostoli, E., Jones, A. P., Pearson, N., Foley, L., Biddle, S. J. H., & Atkin, A. J. (2022). The Association of Contemporary Screen Behaviours with Physical Activity, Sedentary Behaviour and Sleep in Adolescents: a Cross-sectional Analysis of the Millennium Cohort Study. International Journal of Behavioral Medicine.

71. Da silva, I. C. M., Van hees, V. T., Ramires, V. V., Knuth, A. G., Bielemann, R. M., Ekelund, U., Brage, S., & Hallal, P. C. (2014). Physical activity levels in three Brazilian birth cohorts as assessed with raw triaxial wrist accelerometry. International Journal of Epidemiology, 43(6), 1959–1968.

72. van Hees, V. T., Sabia, S., Jones, S. E., Wood, A. R., Anderson, K. N., Kivimäki, M., Frayling, T. M., Pack, A. I., Bucan, M., Trenell, M. I., Mazzotti, D. R., Gehrman, P. R., Singh-Manoux, B. A., & Weedon, M. N. (2018). Estimating sleep parameters using an accelerometer without sleep diary. Scientific Reports, 8(1).

73. Komanduri, S., Jadhao, Y., Guduru, S. S., Cheriyath, P., & Wert, Y. (2017). Prevalence and risk factors of heart failure in the USA: NHANES 2013 – 2014 epidemiological follow-up study. Journal of Community Hospital Internal Medicine Perspectives, 7(1), 15–20.

74. Connelly, R., & Platt, L. (2014). Cohort Profile: UK Millennium Cohort Study (MCS). International Journal of Epidemiology, 43(6), 1719–1725.

75. Millennium Cohort Study Sixth Sweep (MCS6) Prepared for the Centre for Longitudinal Studies, UCL Institute of Education Ipsos MORI | Millennium Cohort Study Sixth Sweep-Technical Report. (2017). http://www.ipsos-mori.com/terms.http://www.ipsos-mori.com/terms.

76. Bates, D., Mächler, M., Bolker, B., & Walker, S. (2015). Fitting Linear Mixed-Effects Models Using lme4. Journal of Statistical Software, 67, 1–48.

77. John, D., Tang, Q., Albinali, F., & Intille, S. (2019). An Open-Source Monitor-Independent Movement Summary for Accelerometer Data Processing. Journal for the Measurement of Physical Behaviour, 2(4), 268–281.

78. Karas, M., Muschelli, J., Leroux, A., Urbanek, J. K., Wanigatunga, A. A., Bai, J., Crainiceanu, C. M., & Schrack, J. A. (2022). Comparison of Accelerometry-Based Measures of Physical Activity: Retrospective Observational Data Analysis Study. JMIR mHealth and uHealth, 10(7), e38077.

79. White, T., Westgate, K., Wareham, N. J., & Brage, S. (2016). Estimation of Physical Activity Energy Expenditure during Free-Living from Wrist Accelerometry in UK Adults. PLOS ONE, 11(12), e0167472.

