## Supplementary Data for "Patterns of physical activity in hunter-gatherer children compared with US and UK children"

### *Supplemental Tables and Figures*

***Table S 1: Summary Statistics presented for BaYaka boys and girls. Reported as mean and standard deviation in brackets, to 2 significant figures.***

| | Overall | Boys | Girls |
|---|---|---|---|
| **n** | 51 | 29 | 22 |
| **Age** | 10.90 (4.35) | 10.20 (4.60) | 11.81 (3.93) |
| **Recorded in 2022** | 23 (45.1) | 12 (41.4) | 11 (50.0) |
| **Mean Acceleration (mg ENMO)** | 46.60 (13.23) | 48.12 (15.45) | 44.61 (9.55) |
| **Sedentary (Min/Day)** | 1066.63 (89.69) | 1071.05 (99.89) | 1060.79 (76.06) |
| **Of Which: Wakeful (Min/Day)** | 585.87 (129.17) | 583.07 (133.14) | 589.55 (126.74) |
| **Light (Min/Day)** | 182.74 (38.38) | 174.24 (38.69) | 193.94 (35.78) |
| **MVPA (Min/Day)** | 199.95 (70.94) | 206.24 (75.14) | 191.66 (65.79) |
| **Of Which: Moderate (Min/Day)** | 179.90 (60.45) | 183.45 (64.96) | 175.21 (55.06) |
| **Of Which: Vigorous (Min/Day)** | 19.36 (14.82) | 21.64 (17.10) | 16.36 (10.79) |
| **Wake-up Time (hh:mm)** | 05:49 (01:05) | 05:46 (01:17) | 05:53 (00:52) |
| **Sleep Onset (hh:mm)** | 21:46 (01:09) | 21:33 (1:24) | 22:01 (00:48) |

***Table S 2: Linear mixed effect model coefficients for mean acceleration and each activity intensity examining the effect of age and gender as fixed effects with and without an interaction term.***

| | **Mean Acceleration** | | **MVPA** | | **Light Activity** | | **Sedentary Activity** | | **Of Which: Wakeful** | |
|---|---|---|---|---|---|---|---|---|---|---|
| ***Predictors*** | *Additive* | *Interaction* | *Additive* | *Interaction* | *Additive* | *Interaction* | *Additive* | *Interaction* | *Additive* | *Interaction* |
| **(Intercept)** | **37.3 *** (27.4 – 47.2)** | **37.7 *** (25.9 – 49.5)** | **119.5 *** (72.2 – 166.9)** | **125.3 *** (69.1 – 181.5)** | **146.2 *** (116.8 – 175.6)** | **148.3 *** (113.3 – 183.3)** | **1167.9 *** (1104.9 – 1230.9)** | **1163.5 *** (1088.6 – 1238.4)** | **647.7 *** (546.2 – 749.3)** | **601.9 *** (486.4 – 717.4)** |
| **Age** | **0.9 * (0.1 – 1.6)** | **0.8 (-0.1 – 1.8)** | **7.9 *** (4.2 – 11.7)** | **7.4 ** (2.8 – 12.0)** | **2.5 * (0.2 – 4.8)** | 2.3 (-0.6 – 5.2) | **-9.9 *** (-14.8 – -5.0)** | **-9.5 ** (-15.7 – -3.4)** | -5.4 (-13.4 – 2.7) | -1.2 (-10.8 – 8.3) |
| **Gender [Girl]** | -4.5 (-11.1 – 2.1) | -5.6 (-25.4 – 14.1) | -24.1 (-56.1 – 8.0) | -41.2 (-136.7 – 54.2) | 14.3 (-5.5 – 34.1) | 8.1 (-50.6 – 66.7) | 7.6 (-34.8 – 49.9) | 20.5 (-105.9 – 146.8) | 30 (-39.0 – 99.0) | 176 (-23.3 – 375.4) |
| **Year [2018]** | 4.6 (-2.0 – 11.1) | 4.5 (-2.1 – 11.2) | 9.5 (-23.0 – 42.0) | 8.9 (-23.8 – 41.7) | 5.5 (-14.1 – 25.0) | 5.3 (-14.6 – 25.1) | -6.7 (-48.7 – 35.4) | -6.3 (-48.8 – 36.3) | -27 (-99.5 – 45.5) | -21.8 (-92.9 – 49.3) |
| **Age × Gender [Girl]** | | 0.3 (-1.3 – 1.9) | | 2.2 (-6.1 – 10.4) | | 2.5 (-2.3 – 7.3) | | -4.9 (-15.4 – 5.6) | | -15.4 (-32.8 – 2.0) |
| **Random Effects** | | | | | | | | | | |
| **$\sigma^2$** | 82.13 | 82.03 | 3231.56 | 3247.39 | 699.95 | 699.82 | 4090.25 | 4094.81 | 21763.38 | 22115.31 |
| **$\tau_{00}$** | 95.31 ID | 98.64 ID | 1685.27 ID | 1704.61 ID | 847.31 ID | 872.91 ID | 3577.49 ID | 3677.25 ID | 5575.15 ID | 4579.81 ID |
| **ICC** | 0.54 | 0.55 | 0.34 | 0.34 | 0.55 | 0.56 | 0.47 | 0.47 | 0.2 | 0.17 |
| **N** | 51 ID | 51 ID | 51 ID | 51 ID | 51 ID | 51 ID | 51 ID | 51 ID | 51 ID | 51 ID |
| **Observations** | 150 | 150 | 147 | 147 | 145 | 145 | 148 | 148 | 145 | 145 |
| **Marginal $R^2$ / Conditional $R^2$** | 0.099 / 0.583 | 0.097 / 0.590 | 0.193 / 0.469 | 0.192 / 0.470 | 0.108 / 0.597 | 0.107 / 0.603 | 0.186 / 0.566 | 0.183 / 0.570 | 0.027 / 0.225 | 0.050 / 0.213 |

***Note: All complete days were included for all individuals. Reference was a Boy recorded in 2022. Values are estimate and 95% confidence interval. *** $p < 0.001$; ** $p < 0.01$; * $p < 0.05$.***

***Table S 3: Linear model estimates for standardised (z-scored) mean activity, adjusting for study, age (Z-Scored) and gender.***

| *Predictors* | **Standardised Mean Activity (Z-Scored)** *Estimates (CI)* |
|---|---|
| **(Intercept)** | 0.06 (-0.18 – 0.29) |
| **Gender [Girl]** | 0.05 (0.00 – 0.11) |
| **Age [Z-Scored]** | 0.22 (-0.01 – 0.46) |
| **Population [NHANES]** | -0.03 (-0.26 – 0.21) |
| **Age × Population [NHANES]** | -0.51 *** (-0.75 – -0.27) |
| Random Effects | |
| **$\sigma^2$** | 0.33 |
| **$\tau_{00}$** | 0.56 id |
| **ICC** | 0.63 |
| **N** | 2772 id |
| **Observations** | 14451 |
| **$R^2$ / $R^2$ adjusted** | 0.082 / 0.658 |

***Note: Reference is a BaYaka boy. Values are estimate and 95% confidence interval * p<0.05 ** p<0.01 *** p<0.001***

***Table S 4: Mixed effect model estimates of wake up and sleep onset times.***

| *Predictors* | **Wake Up Time**<br>*Estimates*<br>*(CI)* | **Sleep Onset**<br>*Estimates*<br>*(CI)* |
|---|---|---|
| **(Intercept)** | 05:00 ***<br>(03:18 – 06:49) | 19:43 ***<br>(17:58 – 21:27) |
| **Age** | 00:04<br>(-00:05 – 00:13) | 00:10 *<br>(00:02 – 0:19) |
| **Gender [Girl]** | 00:05<br>(-01:05 – 01:14) | 00:23<br>(-00:44 – 01:29) |
| | Random Effects | |
| **$\sigma^2$** | 9.79 | 8.96 |
| **$\tau_{00}$ (Individual ID)** | 0.00 $_{ID}$ | 0.00 $_{ID}$ |
| **N** | 23 $_{ID}$ | 23 $_{ID}$ |
| **Observations** | 115 | 115 |
| **Marginal $R^2$ / Conditional $R^2$** | 0.008 / NA | 0.051 / NA |

***Note: Estimates are presented as hh:mm. Intercepts represent a time of day, with effects showing a deviation in hours and minutes. The effect of age and gender are included as fixed effects with the day of recording nested within each individual as a random effect. All complete days were included for all individuals from the 2022 data collection. Reference is a Boy aged 0. Values are estimate and 95% confidence interval. *** p < 0.001; ** p < 0.01; * p < 0.05.***

***Table S 5: Linear mixed effect model coefficients for the mean acceleration at each three-hour interval of time.***

| | **Mean Acceleration (mg ENMO)** |
|---|---|
| *Predictors* | *Estimates (CI)* |
| **(Intercept)** | -5.47 (-16.85 – 5.91) |
| **03:00 – 06:00** | -1.14 (-6.37 – 4.08) |
| **06:00 – 09:00** | 41.17 *** (35.94 – 46.39) |
| **09:00 – 12:00** | 83.30 *** (78.07 – 88.52) |
| **12:00 – 15:00** | 65.30 *** (60.08 – 70.52) |
| **15:00 – 18:00** | 65.53 *** (60.31 – 70.76) |
| **18:00 - 21:00** | 42.56 *** (37.34 – 47.78) |
| **21:00 – 24:00** | 5.73 * (0.51 – 10.95) |
| **Age** | 1.31 ** (0.37 – 2.24) |
| **Gender [Girl]** | -4.52 (-11.71 – 2.67) |
| **Random Effects** | |
| **$\sigma^2$** | 407.30 |
| **$\tau_{00}$ (weekday:ID)** | 20.37 |
| **$\tau_{00}$ (ID)** | 59.59 |
| **ICC** | 0.16 |
| **$N_{weekday}$** | 5 |
| **$N_{ID}$** | 23 |
| Observations | 920 |
| Marginal $R^2$ / Conditional $R^2$ | 0.667 / 0.722 |

***Note: The effect of age and gender are included as fixed effects with the day of recording nested within each individual as a random effect. All complete days were included for all individuals. Reference is a Boy aged 0. Values are estimate and 95% confidence interval. *** p < 0.001; ** p < 0.01; * p < 0.05.***

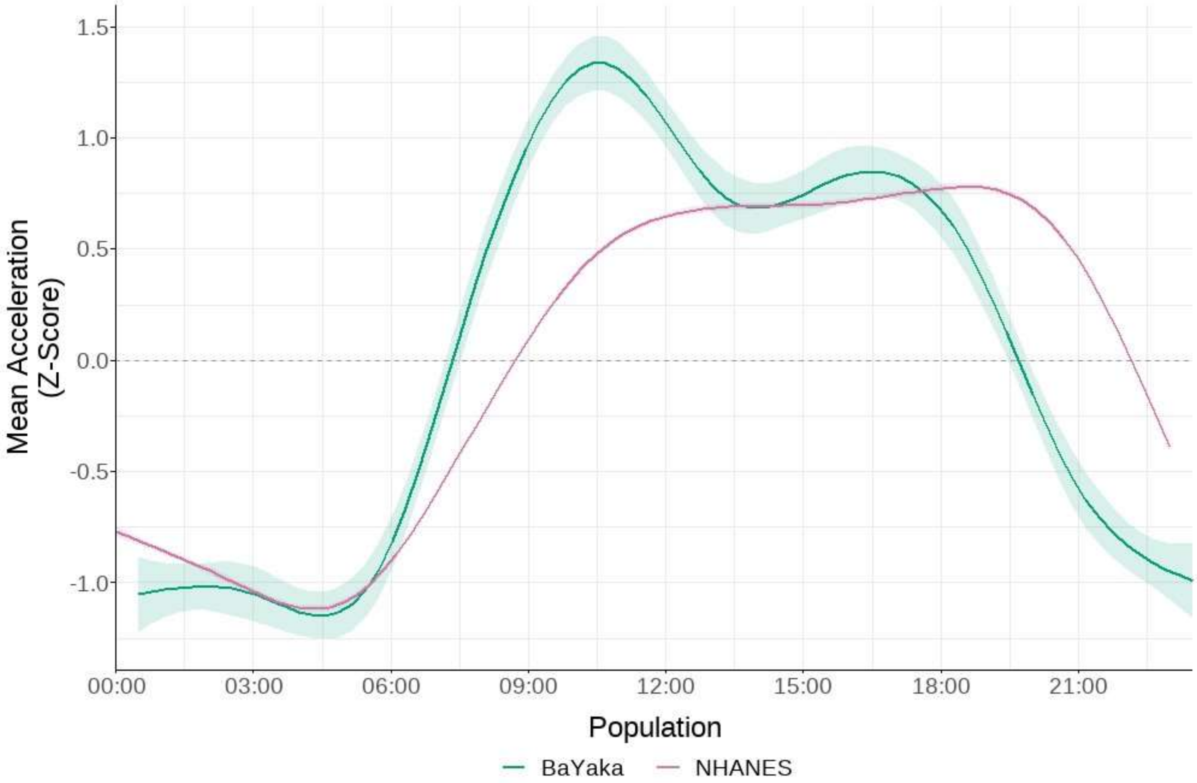


***Figure S 1: Distribution of Z-Scored physical activity throughout the day in the BaYaka and NHANES datasets. Accelerations were Measures of acceleration are standardised within each sample. Sample includes all children in the BaYaka and NHANES with 5 complete days of recording. For all individuals their mean acceleration at each hour is averaged across the 5 days of recording. The mean acceleration was calculated for each hour interval and presented at the half hour (i.e., the mean acceleration between 09:00 and 10:00 is presented at 09:30). Z-Scored accelerations are used to account for the different processing techniques employed in the BaYaka and NHANES studies.***

***Table S 6: Mixed effect model of standardised (Z-Scored) mean accelerations in three hour windows across the day in the BaYaka and NHANES sample.***

| | **Mean Acceleration (Z-Score)** | **NHANES:Mean Acceleration (Z-Score)** |
|---|---|---|
| *Predictors* | *Estimates* | |
| **00:00 – 03:00** | 0.03 (-0.14 – 0.20) | 0.02 (-0.14 – 0.19) |
| **03:00 – 06:00** | -0.51 *** (-0.70 – -0.32) | 0.11 (-0.07 – 0.30) |
| **06:00 – 09:00** | 1.11 *** (0.94 – 1.28) | -0.35 *** (-0.52 – -0.18) |
| **09:00 – 12:00** | 2.22 *** (2.05 – 2.38) | -0.83 *** (-1.00 – -0.66) |
| **12:00 – 15:00** | 1.74 *** (1.58 – 1.91) | -0.16 (-0.33 – 0.01) |
| **15:00 – 18:00** | 1.75 *** (1.58 – 1.92) | -0.12 (-0.29 – 0.05) |
| **18:00 - 21:00** | 1.15 *** (0.98 – 1.31) | 0.37 *** (0.20 – 0.54) |
| **21:00 – 24:00** | 0.18 * (0.01 – 0.35) | 0.44 *** (0.27 – 0.61) |
| **Gender [Girl]** | -0.02 (-0.04 – 0.01) | |
| **Age** | -0.05 *** (-0.05 – -0.04) | |
| **Random Effects** | | |
| **$\sigma^2$** | 0.42 | |
| **$\tau_{00\ weekday:id}$** | 0.03 | |
| **$\tau_{00\ id}$** | 0.11 | |
| **ICC** | 0.25 | |
| **$N_{weekday}$** | 12 | |
| **$N_{id}$** | 2483 | |
| Observations | 135925 | |
| Marginal $R^2$ / Conditional $R^2$ | 0.438 / 0.579 | |

***Note: Accelerations were averaged over three hour windows and z-scored to account for different means of presenting accelerometer outputs (See Table S 9). The effect of age and gender are included as fixed effects with the day of recording nested within each individual as a random effect. All complete days were included for all individuals. Reference is the standardised acceleration of a BaYaka child (Left Column) between 3am and 6am. Right hand column represents difference between NHANES and BaYaka in a given time window. *** p < 0.001; ** p < 0.01; * p < 0.05.***

***Table S 7: Intercept only mixed effect models of Z-Scores physical Activity in the BaYaka and NHANES sample.***

| | **BaYaka (Z-Scored Mean Acceleration)** | **NHANES (Z-Scored Mean Acceleration)** |
|---|---|---|
| | *Estimates (CI)* | *Estimates (CI)* |
| *(Intercept)* | -0.00 (-0.32 – 0.32) | 0.07 *** (0.04 – 0.10) |
| ***Random Effects*** | | |
| ***$\sigma^2$*** | 0.51 | 0.33 |
| ***$\tau_{00}$*** | 0.51 ID | 0.64 ID |
| ***ICC*** | 0.50 | 0.66 |
| ***N*** | 23 ID | 2721 ID |
| ***Observations*** | 115 | 14301 |
| ***Marginal $R^2$ / Conditional $R^2$*** | 0.000 / 0.497 | 0.000 / 0.657 |

***Note: Each individual ID is included as a random effect. All complete days were included for all individuals from the 2022 data collection in the BaYaka and the 2013 NHANES sample. Values are estimate and 95% confidence interval. *** p < 0.001; ** p < 0.01; * p < 0.05.***

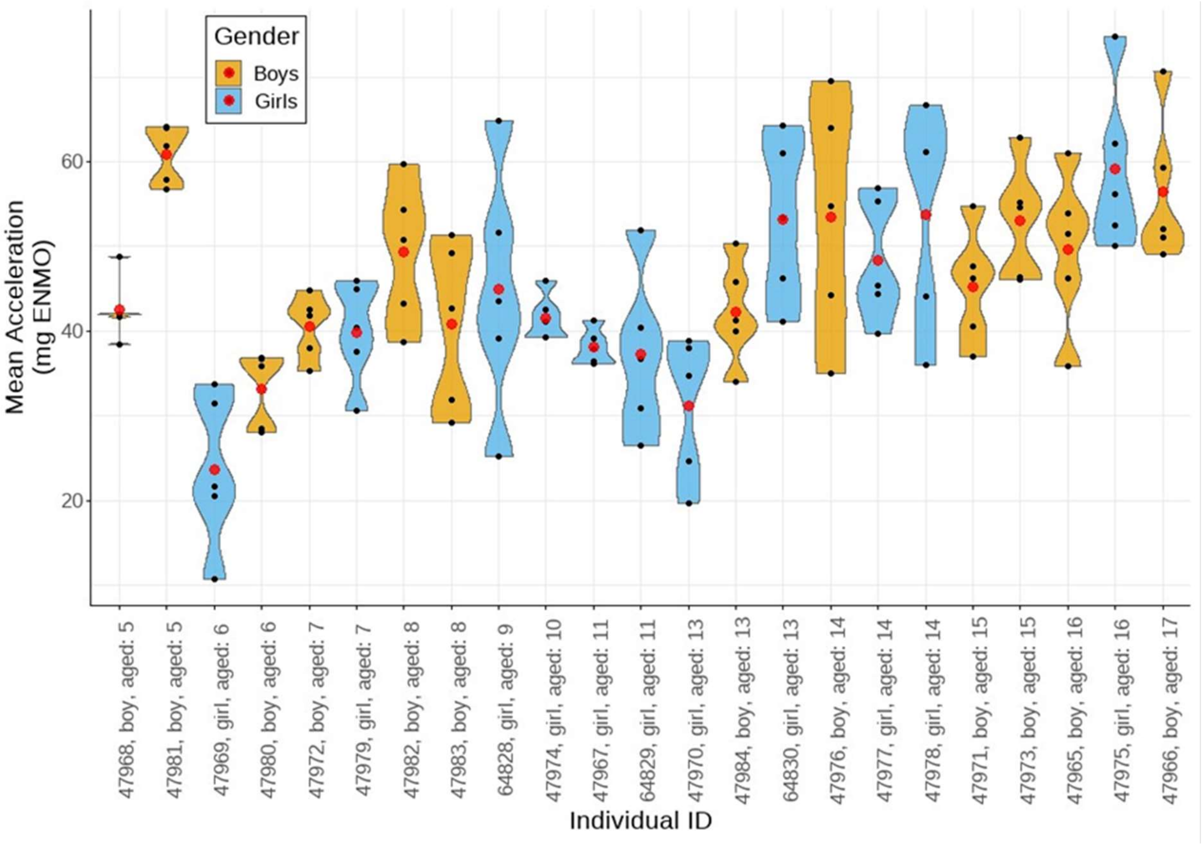


***Figure S 2: Violin plot of individual variation in mean acceleration across 5 days of recording in the 2022 field trip. Red point represents each individuals mean acceleration across all complete days of recording, each grey point represents the mean acceleration on a given day of recording. Individuals are arranged in increasing age, with colours set by gender.***

***Table S 8: Millennium Cohort Summary Data. Data presented comes from the 6<sup>th</sup> sweep of the Millennium Cohort when participants were aged 14.***

| | Overall | Boys | Girls |
|---|---|---|---|
| **n** | 4533 | 2182 | 2351 |
| **Age (Years)** | 14.23 (0.50) | 14.23 (0.56) | 14.23 (0.44) |
| **Height (cm)** | 163.86 (8.07) | 166.68 (8.70) | 161.23 (6.41) |
| **Weight (kg)** | 57.55 (12.80) | 57.90 (13.40) | 57.22 (12.18) |
| **BMI** | 21.34 (4.07) | 20.70 (3.84) | 21.95 (4.19) |
| **Body Fat (%)** | 21.91 (9.13) | 16.46 (7.89) | 27.09 (6.97) |
| **Season (%)** | | | |
| **Autumn** | 961 (22.0) | 472 (22.6) | 489 (21.5) |
| **Spring** | 1410 (32.3) | 671 (32.1) | 739 (32.5) |
| **Summer** | 1385 (31.7) | 651 (31.1) | 734 (32.3) |
| **Winter** | 608 (13.9) | 297 (14.2) | 311 (13.7) |
| **Ethnicity (%)** | | | |
| **White** | 3714 (83.1) | 1785 (83.0) | 1929 (83.1) |
| **Black or Black British** | 116 ( 2.6) | 53 ( 2.5) | 63 ( 2.7) |
| **Indian** | 112 ( 2.5) | 60 ( 2.8) | 52 ( 2.2) |
| **Pakistani or Bangladeshi** | 251 ( 5.6) | 109 ( 5.1) | 142 ( 6.1) |
| **Mixed** | 181 ( 4.0) | 97 ( 4.5) | 84 ( 3.6) |
| **Other** | 97 ( 2.2) | 46 ( 2.1) | 51 ( 2.2) |
| **Mean Acceleration (Mg ENMO)** | 34.01 (13.26) | 36.81[a] (15.31) | 31.42[a] (10.36) |
| **MVPA (mean (SD))** | 128.58 (54.86) | 134.10[b] (59.77) | 123.46[b] (49.34) |

***Note: Further information on the study can be found at " https://cls.ucl.ac.uk/cls-studies/millennium-cohort-study/ ". Data is accessible via the UK Data Service. a = in linear models adjusted for age and BMI, compared to boys the mean acceleration amongst girls was lower (β = -5.10, SE = 0.40, p <0.001). b = in linear models adjusted for age and BMI, compared to boys the volume of MVPA amongst girls was lower (β= -10.05, SE = 1.67, p <0.001).***

***Table S 9: NHANES summary data. Data used in the present study comes from the 2013-14 data collection of the National Health and Nutrition Exmaination Survey.***

| | Overall | Boys | Girls |
|---|---|---|---|
| **n** | 3308 | 1699 | 1609 |
| **Age (mean (SD))** | 10.65 (4.55) | 10.55 (4.55) | 10.76 (4.56) |
| **Ethnicity** | | | |
| **Mexican American** | 729 (22.0) | 350 (20.6) | 379 (23.6) |
| **Other Hispanic** | 333 (10.1) | 176 (10.4) | 157 ( 9.8) |
| **Non-Hispanic White** | 854 (25.8) | 462 (27.2) | 392 (24.4) |
| **Non-Hispanic Black** | 844 (25.5) | 441 (26.0) | 403 (25.0) |
| **Other** | 548 (16.6) | 270 (15.9) | 278 (17.3) |
| **Mean Acceleration (MIMS Unit)* (mean (SD))** | 14711.23 (6051.93) | 14563.69[c] (6220.36) | 14860.63[c] (5874.98) |

***Note: *MIMS: Monitor Independent Motion Sensing. A measure of total acceleration experienced by the device, averaged between days. More information on the NHANES dataset and access can be found at " https://wwwn.cdc.gov/nchs/nhanes/Default.aspx ". c = In linear models adjusted for age, girls and boys recorded similar MIMS-Units per day (β = -209, SE = 248.52, p = 0.40).***

A limitation to NHANES is that while the devices are broadly similar (triaxial wrist-worn accelerometer), thus capturing the same latent variable, the means of processing the data varies from those employed with the BaYaka. NHANES employs a metric called MIMS (Monitor Independent Motion Sensing)[77], a technique similar to HPFVM (High-Pass Filter Vector Magnitude).[78] The difference between this and mg ENMO, is that mg ENMO treats gravity as a constant, meaning 1g is subtracted from the acceleration; a HPFVM or MIMS technique does not assume gravity is constant and instead treats gravity as a low frequency component to be filtered out.[79] While some estimates of a conversion factor have been made,[77] these have not been validated. Accordingly, comparisons with NHANES examine trends in patterns of activity using internally estimated z-scored volumes of activity. Using standardised activity measures precludes the ability to compare total volumes, but the study design does allow for comparisons of how activity varies across ages and within days.

The same process of standardisation was undertaken to examine difference between populations in the distribution of activity throughout the day. As before, activity was averaged over three-hour windows. Mixed effect linear models were used to model the effect of time of day on proportional physical activity, including the study population as an interaction term for time of day, with age and gender included as covariates. Individual ID and day of the week were included as random effects. The reference time window in this model differs from the analysis of activity across the day in the BaYaka. to improve the interpretability of the output the time window of 03:00-06:00 was used as the reference as difference between populations was at a minimum during this time window.
